# C3PI: Component Puzzle Protein-Protein Interaction Prediction

**DOI:** 10.1101/2025.07.26.666948

**Authors:** SeyedMohsen Hosseini, G. Brian Golding, Lucian Ilie

## Abstract

Proteins primarily perform their functions through interactions with other proteins, making the accurate prediction of protein-protein interactions (PPIs) a fundamental problem. Experimental methods for determining PPIs are often slow and expensive, which has driven significant efforts to improve the performance of computational methods in this field. While many methods have been designed, recent thorough investigations proved that the existing methods learn exclusively from sequence similarities and node degrees. When such data leakage is avoided, performances were shown to become random. We introduce C3PI, a novel sequence-based deep learning framework designed for predicting PPIs. C3PI uses as input ProtT5 protein embeddings into a complex architecture that includes two novel components, a puzzler and an entangler, which significantly enhance the model’s performance. Through extensive comparisons with state-of-the-art methods across many datasets, C3PI consistently outperforms competing approaches, especially in key metrics such as AUPRC and AUROC. Most importantly, C3PI is the first PPI prediction method to achieve a significant improvement over random on the leakage-free gold standard dataset. C3PI is available as a web server at c3pi.csd.uwo.ca and source code from github.com/lucian-ilie/C3PI.

## Introduction

Proteins play a crucial role in various cellular processes, including cell growth, gene expression, and intercellular communication [2]. In the late 1990s, analyses of protein functions predominantly focused on individual proteins [15]. However, to gain a comprehensive understanding of protein functionality, it is essential to study proteins in relation to their interacting partners. Since proteins operate collaboratively to ensure proper functionality, examining them within the context of their interactions is imperative. With the publication of the human genome [33] and advancements in proteomics, it has become increasingly important to understand not only the functions of individual proteins but also how they interact with one another within cellular environments.

Protein-protein interactions (PPIs) play numerous critical roles in our bodies, serving as a foundation for various biological processes. One significant application of understanding PPIs is in predicting the function of target proteins, which can provide insights into their roles within the cellular environment. Additionally, PPIs are crucial for determining the effectiveness of drugs, as many therapeutic agents function by modulating these interactions to achieve a desired biological response [24].

In the context of a living organism or cell, PPIs are interpreted as physical contacts between proteins, which occur through molecular docking mechanisms. These interactions enable proteins to form complexes, communicate signals, and execute a wide range of cellular functions essential for maintaining life [11]. Understanding the specifics of how proteins interact at a molecular level not only enhances our knowledge of fundamental biological processes but also aids in the development of new therapeutic strategies and drug designs.

Experimental methods for identifying protein interactions, such as immunoprecipitation utilizing protein-specific antibodies and Protein A/G affinity beads, are known for being time-consuming, labor-intensive, and expensive [21]. Coimmunoprecipitation is widely recognized as a standard technique in the field, employing antibodies and affinity beads to identify molecules that interact with proteins. Another experimental approach is the pull-down assay, which involves affinity purification with multiple wash and elution steps [22]. Additionally, techniques like surface plasmon resonance (SPR) [12], bacterial two-hybrid [18], and cytology two-hybrid [4] are used to investigate protein interactions in vitro.

In contrast to experimental methods, computational approaches are rapid and cost-effective. Consequently, numerous studies aim to predict protein interactions using computational methods. The protein interaction prediction problem is typically framed as a classification challenge. Classifiers undergo two key phases: training, where the classifier is educated on available data, and testing, where the classifier is tasked with predicting the correct class for input data. Given the potential for prediction errors, multiple metrics are employed to assess the performance of classifiers.

We focus in this study on sequence-based programs for two reasons. First, the number of available protein sequences outnumbers that of structures by two orders of magnitude [23]. Second, while the advent of AlphaFold [17, 1] helped predict significantly better structures than before, it cannot reliably predict structures for proteins containing intrinsically disordered regions, which are the ones having the most interactions. Therefore, sequence-based prediction is still the best solution. Many sequence-based programs have been designed for PPI prediction, employing a variety of methods. Most popular ones include DeepFE [30], DPPI [16], R-FC and R-LSTM [25], Bio2Vec [34], PIPR [9], StackPPI D-SCRIPT [28], TAGPPI [29], and Topsy-Turvy [27].

The performance of the computational methods has often been claimed to have very high accuracy (95-99%) [5], seemingly implying that this is a solved problem. Bernett *et al*. [5] noticed that the overlaps between training and test sets resulting from random splitting lead to strongly overestimated performances. They investigated this issue thoroughly and their conclusions about the PPI prediction problem is that not only it is not solved, but it is actually wide open. They prove that the current models learn exclusively from sequence similarity and node degree (number of interacting proteins), rather than identifying more complex sequence features representing binding pockets, protein domains or similar motifs [5]. When data leakage is avoided, performances were shown to become random. This render the methods unequipped to handle interactions of dark proteins [7], which have no known similarity with other sequences. In order to help future development of sequence-based PPI prediction methods, they built a *gold standard* dataset, with training, validation and testing components, which is free from data leakage. They have tested many programs on this dataset and proved that their performance is random or very close.

In this paper we introduce C3PI, a new deep learning program for PPI prediction. C3PI uses as input ProtT5 protein embeddings [13] into a complex architecture that includes two novel components, a puzzler and an entangler. The puzzler produces random permutations of segments of each input protein sequence, whereas the entangler performs a mid-level combination of features extracted from the two input sequences. We perform a comprehensive evaluation of its performance by comparing it with state-of-the-art tools on six species datasets used by top competitors, as well as on the gold standard dataset mentioned above. C3PI has the highest AUROC and AUPRC parameters in all but one species tests. On the gold standard dataset, C3PI is the first method to achieve significant improvement over random.

## Material and Methods

### Datasets

To assess the performance of C3PI in predicting PPIs, we utilized two types of datasets: the *species* datasets used by D-SCRIPT [31] and the *gold standard* dataset developed by Bernett *et al*. [5].

The species datasets have been sourced from the STRING database (version 11) [32] and processed to keep only high-confidence interactions, as well as filtered out by clustering proteins at a 40% similarity threshold using CD-HIT [20, 14], in order to mitigate the model’s reliance on sequence similarity alone for memorizing interactions. Negative PPIs have been produced by randomly pairing proteins from the non-redundant set [16].

Our human PPI dataset follows different guidelines. While D-SCRIPT maintains a 10:1 ratio for positive and negative interactions, we opted for a 1:1 ratio in the training set and 10:1 in the validation set. This was done to achieve better training while maintaining universal applicability to many species. The fairly large total number of pairs allowed us to employ about 95% of the data for training, and the remaining for validation, trying to maximize the amount of training data while preserving a meaningful validation set.

The gold standard dataset is taken from Bernett *et al*. [5]. It was purposefully built to be free from data leakage and minimized pairwise sequence similarities. We refer to [5] for details. All datasets are shown in Table 1.

**Table 1.** Datasets.

| Species datasets | Pairs | Proteins | Lengths | Avg. len. |
| --- | --- | --- | --- | --- |
| *E.coli* | 22,000 | 4,437 | 50 – 799 | 286.42 |
| *S.cerevisiae* | 55,000 | 5,664 | 50 – 800 | 341.53 |
| *D.melanogaster* | 55,000 | 19,213 | 50 – 800 | 387.97 |
| *C.elegans* | 55,000 | 25,429 | 50 – 800 | 351.70 |
| *M.musculus* | 55,000 | 37,497 | 50 – 800 | 392.85 |
| *H.sapiens* (train) | 95,864 | 14,272 | 50 – 800 | 382.79 |
| *H.sapiens* (validate) | 4,219 | 1,536 | 51 – 800 | 381.38 |
| *H.sapiens* (test) | 52,725 | 15,525 | 50 – 800 | 382.65 |
| Gold standard dataset (*H.sapiens*) |  |  |  |  |
| Train (INTRA1) | 163,192 | 4,286 | 54 – 14,507 | 752.71 |
| Validate (INTRA0) | 59,260 | 3,711 | 56 – 34,350 | 590.48 |
| Test (INTRA2) | 52,048 | 3,022 | 51 – 5,183 | 504.73 |

### Architecture

The proposed PPI prediction model is organized as a two branch, multi-scale feature extractor followed by a fusion and classification head. Each branch processes one protein sequence ProtT5 embedding independently, extracting features at six distinct length scales. The outputs from both branches are then fused, aggregated into a single representation, and passed through a final multilayer classifier to calculate probability of interactions.

#### Puzzler

Protein embeddings do not directly encode three dimensional proximity of nonadjacent residues. It is often the case that residues that are far away in the sequence become spatially close in the three dimensional structure. Therefore, the main idea of the *puzzler* component is to expose the model to alternative residue ordering. Each protein sequence is divided into pieces which are then shuffled, to produce a *permuted* sequence. These permuted sequences are then fed as input to the model. Precisely, each protein sequence is first normalized to a length of 800, as customary in the area, by either padding or truncating, and then split into 15 pieces of size 53 each; this uses the first 795 residues, while the last 5 are ignored. These pieces are randomly shuffled by the puzzler to produce 8 permuted sequences for each input protein sequence; see Figure 1. Each input protein is permuted according to 8 fixed permutations, the first of which is the identity, which means the original sequence is always used. During training, for each input protein pair, the puzzler generates 8 pairs by combining the corresponding permutations. Each of these is treated as a separate input pair with the same property as the original one. Each pair is passed through the model individually, allowing it to learn from varied representations of the same underlying data. At inference, 8 separate permuted pairs are again generated using the same permutations and each instance is independently passed through the trained model to produce a prediction. The final output is obtained by averaging the 8 predictions.

**Fig. 1.**
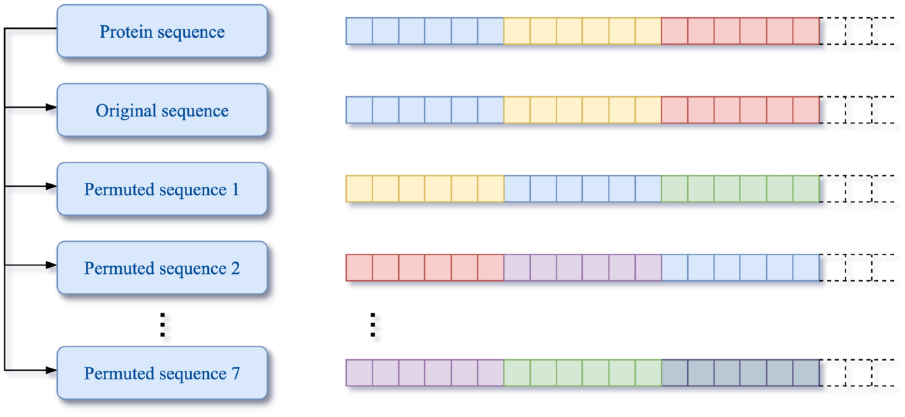
The puzzler component. The original sequence produces 8 permuted versions, the first of which is the original sequence.

Different sets of random permutations have been tested but no significant difference in predicting power over the training dataset has been detected. While the idea deserves further analysis, it appears that as long as parts of the protein sequences are moved around, the prediction power improves significantly. The Ablation section indicates the power of the puzzler component of the architecture.

Also, limited testing has been performed for the size of the permuted pieces. For example, splitting each sequence of length 800 into the more natural 16 blocks of length 50 produces a model that is slightly less effective. This, like other aspects of the architecture, could potentially benefit from further tuning.

#### Multi-scale Convolutional Dense Block

At the core of each branch lies a set of six convolutional dense modules, designed to capture sequence patterns ranging from global to highly local. Each module applies a one dimensional convolution with a specific kernel size and stride, followed by batch normalization and a nonlinear activation. The resulting feature maps are then flattened and projected via a fully connected layer into a fixed dimensional embedding. A dropout layer follows each dense projection to reduce overfitting.

The six (kernel, stride, output-dim) configurations are chosen as: (795, 1, 16), (400, 200, 32), (200, 100, 64), (100, 50, 128), (50, 25, 256), (20, 10, 512). These settings ensure that the first module effectively sees the entire input sequence (capturing global context), while the last module focuses on short subsequences with a stride of 10 (capturing very local patterns). Flattening and projecting into a relatively low-dimensional embedding at each scale encourages the network to summarize each receptive field into a compact feature vector. Figure 2 shows the overall structure of a convolutional dense block block.

**Fig. 2.**
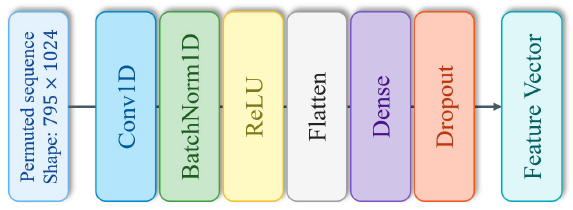
Multi-scale convolutional dense block. Each configuration captures protein sequence features at a specific resolution, ranging from global to highly local patterns.

#### Entangler

To model protein-protein interactions, two parallel branches with identical structures independently process the embeddings of the two input proteins. For each protein sequence, *p* = 1, 2, each branch produces six embeddings 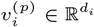 with *d*_*i*_ ∈ {16, 32, 64, 128, 256, 512}. By processing each protein through the same hierarchy of kernel sizes and strides, each branch learns to extract complementary features at matching scales for the two partners. Figure 3 shows the overall structure of a branch block.

**Fig. 3.**
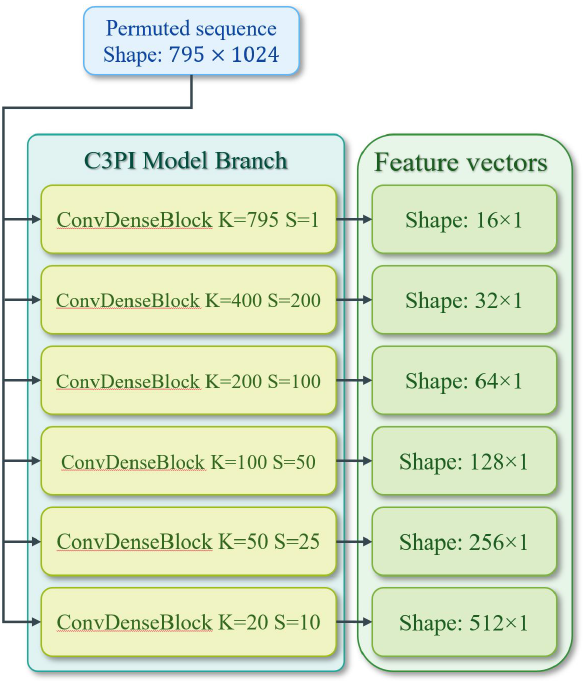
Architecture of a single CNN branch. Each branch processes one protein and extracts six scale-specific embeddings using the multi scale convolutional dense block.

Rather than fusing both protein embeddings only at the very end, the *entangler* performs a mid level combination at each scale. Specifically, for scale *i*, the two feature vectors 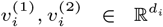 are concatenated into a 2*d*_*i*_ dimensional vector. This concatenated vector is passed through a small fully connected layer, mapping 2*d*_*i*_ → *d*_*i*_, with a nonlinear activation and dropout. The intuition behind this design is that it allows the network to learn interaction between two protein features that are specific to a given length scale and keep each fused vector at the same dimensionality, *d*_*i*_, as its original scale, simplifying downstream aggregation. Figure 4 shows the overall structure of an entangler block.

**Fig. 4.**
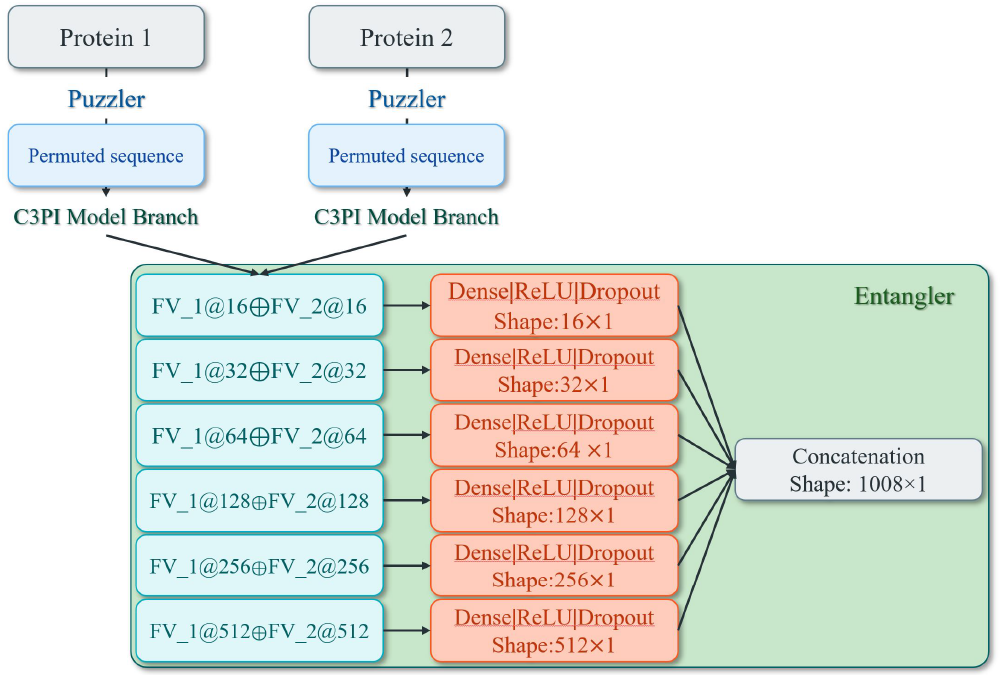
Scale-wise entangler block. At each scale, embeddings from both protein branches are concatenated and passed through a shared transformation to model interactions

#### Aggregation and Final Classification

After generating the six fused vectors, these are concatenated into a single high-dimensional representation of total size 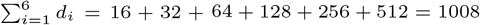.

This 1008-dimensional vector is then passed through a three-layer multilayer perceptron: (i) a linear projection 1008 → 64, followed by ReLU and dropout (rate 0.3); (ii) a linear projection 64 → 8, followed by ReLU and dropout (rate 0.3), and (iii) a linear projection 8 → 1, followed by a sigmoid activation. The final scalar output represents the predicted probability that the two input proteins interact. Figure 5 shows the overall structure of a classifier block.

**Fig. 5.**
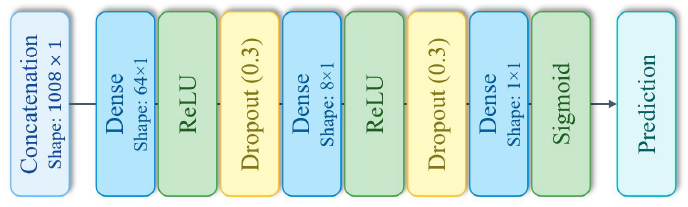
Final classifier block. The six fused embeddings are aggregated and passed through a multi layer perceptron to predict protein protein interaction probability.

## Results

### Competing Methods

When evaluating methods to compare with, one method stood out, Topsy-Turvy [27], which is in agreement with the study of Bernett *et al*. [5]. Therefore, we have spent a considerable amount of effort on this method. We ran both D-SCRIPT, the previous method from the same group, and Topsy-Turvy however, while the D-SCRIPT results match those in their paper, the Topsy-Turvy ones do not. We have communicated with the authors, who helped greatly by providing several trained models, but unfortunately none could match the results in the paper. Therefore, we present both the reported results and the best of the models sent to us. In addition, we include the results from another competitive method, PIPR. All these methods were included also in the study of Bernett *et al*. [5].

### Evaluation scheme

In order to assess the performance of the tested models, we have considered many parameters: sensitivity (recall), specificity, precision, accuracy, F1-score, MCC, and the area under the receiver operating characteristic curve (AUROC) and precision recall curve (AUPRC). The AUPRC is the best indicator of performance for skewed data [10, 26], which makes it most relevant for imbalanced datasets. For balanced datasets, both AUROC and AUPRC reflect well the overall performance of the models.

### Species Data

We compare in this section the performance of C3PI against D-SCRIPT, PIPR, and Topsy-Turvy on the species datasets from Table 1. The results shown in Table 2, with ROC and PR curves shown for the method we run in Figure 6.

**Table 2.** Performance comparisons of methods across six different species using six parameters: sensitivity, specificity, precision, accuracy, F1-score, Matthew’s correlation coefficient, and areas under the ROC and PR curves. The results for the methods marked by * were taken from their papers; the rest were computed by us. The best results are shown in boldface (excluding *).

| Model | Sens. | Spec. | Prec. | Acc. | F1 | MCC | ROC | PR |
| --- | --- | --- | --- | --- | --- | --- | --- | --- |
| *E.coli* |  |  |  |  |  |  |  |  |
| D-SCRIPT* | 0.520 |  | 0.791 |  | 0.627 |  | 0.863 | 0.571 |
| D-SCRIPT | 0.382 | **0.987** | **0.788** | **0.921** | 0.515 | **0.515** | 0.860 | 0.574 |
| PIPR* | 0.131 |  | 0.629 |  | 0.217 |  | 0.675 | 0.308 |
| PIPR+D* | 0.394 |  | 0.793 |  | 0.526 |  | 0.863 | 0.588 |
| Topsy-Turvy* |  |  |  |  |  |  | 0.805 | 0.556 |
| Topsy-Turvy | 0.464 | 0.930 | 0.451 | 0.879 | 0.457 | 0.389 | 0.643 | 0.405 |
| C3PI | **0.675** | 0.910 | 0.429 | 0.889 | **0.524** | 0.475 | **0.873** | **0.633** |
| *S.cerevisiae* |  |  |  |  |  |  |  |  |
| D-SCRIPT* | 0.223 |  | 0.706 |  | 0.339 |  | 0.789 | 0.405 |
| D-SCRIPT | 0.223 | **0.991** | **0.706** | **0.921** | 0.339 | 0.368 | 0.789 | 0.405 |
| PIPR* | 0.085 |  | 0.398 |  | 0.140 |  | 0.718 | 0.230 |
| PIPR+D* | 0.225 |  | 0.708 |  | 0.341 |  | 0.789 | 0.417 |
| Topsy-Turvy* |  |  |  |  |  |  | 0.850 | 0.534 |
| Topsy-Turvy | 0.544 | 0.942 | 0.484 | 0.906 | **0.512** | **0.461** | 0.762 | 0.431 |
| C3PI | **0.690** | 0.893 | 0.391 | 0.874 | 0.499 | 0.456 | **0.886** | **0.536** |
| *D.melanogaster* |  |  |  |  |  |  |  |  |
| D-SCRIPT* | 0.359 |  | 0.798 |  | 0.495 |  | 0.824 | 0.552 |
| D-SCRIPT | 0.359 | **0.991** | **0.798** | 0.933 | 0.495 | 0.508 | 0.824 | 0.552 |
| PIPR* | 0.121 |  | 0.521 |  | 0.196 |  | 0.728 | 0.278 |
| PIPR+D* | 0.361 |  | 0.798 |  | 0.497 |  | 0.824 | 0.562 |
| Topsy-Turvy* |  |  |  |  |  |  | 0.921 | 0.713 |
| Topsy-Turvy | 0.741 | 0.955 | 0.621 | **0.935** | **0.675** | **0.643** | 0.896 | 0.649 |
| C3PI | **0.823** | 0.892 | 0.432 | 0.886 | 0.567 | 0.543 | **0.931** | **0.683** |
| *C.elegans* |  |  |  |  |  |  |  |  |
| D-SCRIPT* | 0.306 |  | 0.840 |  | 0.449 |  | 0.813 | 0.548 |
| D-SCRIPT | 0.306 | **0.994** | **0.840** | 0.932 | 0.448 | 0.482 | 0.813 | 0.548 |
| PIPR* | 0.142 |  | 0.673 |  | 0.235 |  | 0.757 | 0.346 |
| PIPR+D* | 0.308 |  | 0.841 |  | 0.451 |  | 0.814 | 0.559 |
| Topsy-Turvy* |  |  |  |  |  |  | 0.906 | 0.700 |
| Topsy-Turvy | 0.645 | 0.970 | 0.683 | **0.941** | 0.663 | 0.631 | 0.851 | 0.616 |
| C3PI | **0.718** | 0.957 | 0.625 | 0.935 | **0.668** | **0.634** | **0.943** | **0.740** |
| *M.musculus* |  |  |  |  |  |  |  |  |
| D-SCRIPT* | 0.346 |  | 0.818 |  | 0.486 |  | 0.833 | 0.580 |
| D-SCRIPT | 0.346 | **0.992** | **0.818** | **0.934** | 0.486 | 0.506 | 0.833 | 0.580 |
| PIPR* | 0.331 |  | 0.734 |  | 0.456 |  | 0.839 | 0.526 |
| PIPR+D* | 0.355 |  | 0.820 |  | 0.495 |  | 0.838 | 0.609 |
| Topsy-Turvy* |  |  |  |  |  |  | 0.934 | 0.735 |
| Topsy-Turvy | 0.749 | 0.948 | 0.592 | 0.930 | **0.661** | **0.628** | 0.885 | **0.636** |
| C3PI | **0.800** | 0.883 | 0.406 | 0.875 | 0.538 | 0.512 | **0.923** | 0.622 |
| *H.sapiens* |  |  |  |  |  |  |  |  |
| D-SCRIPT* | 0.386 |  | 0.833 |  | 0.527 |  | 0.854 | 0.617 |
| D-SCRIPT | 0.386 | **0.992** | **0.833** | 0.937 | 0.527 | 0.541 | 0.854 | 0.617 |
| PIPR* | 0.701 |  | 0.838 |  | 0.763 |  | 0.960 | 0.835 |
| PIPR+D* | 0.400 |  | 0.949 |  | 0.562 |  | 0.962 | 0.844 |
| Topsy-Turvy | 0.782 | 0.958 | 0.651 | **0.942** | **0.711** | 0.682 | 0.895 | 0.703 |
| C3PI | **0.914** | 0.934 | 0.581 | 0.932 | 0.710 | **0.696** | **0.977** | **0.878** |

**Fig. 6.**
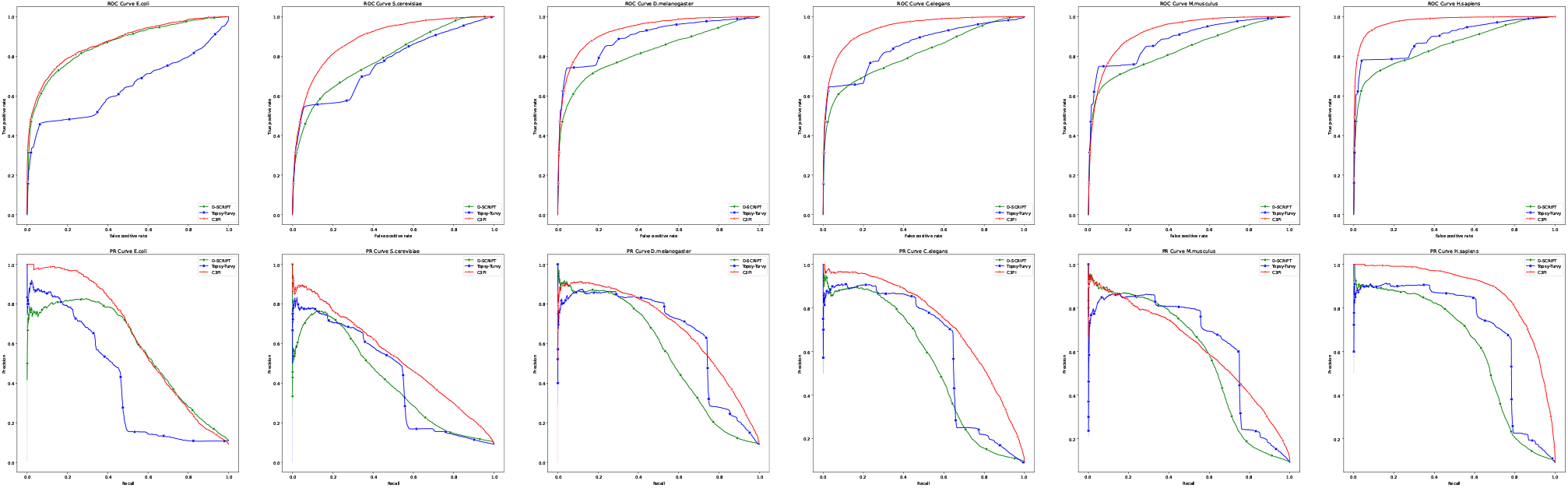
ROC and PR curves for the three models that have been run by us from Table 2: D-SCRIPT, Topsy-Turvy and C3PI for the species datasets.

For the threshold-dependent parameters, the winners are distributed among the methods: sensitivity is won by C3PI in all tests, specificity and precision are won by D-SCRIPT in all tests, accuracy is distributed between D-SCRIPT and Topsy-Turvy, F1-score and MCC are distributed between C3PI and Topsy-Turvy. The most important, threshold-independent, parameters, that measure the entire behaviour of a method on a given dataset, AUROC and AUPRC, are won by C3PI with one exception, the AUPRC for *M*.*musculus*. The second best is Topsy-Turvy with two exceptions, both for *E*.*coli*, where D-SCRIPT is second best.

The average improvement of C3PI over the top competitor, Topsy-Turvy, is 13.35% for AUROC and 26.11% for AUPRC. Given that these are skewed datasets, AUPRC is the most important parameter. In the single case where Topsy-Turvy performs better, it is by 2.09%.

### Gold Standard Dataset

The most relevant comparison is on the gold standard dataset of Bernett *et al*. [5]. Here we compare against all methods they studied (except baseline methods): D-SCRIPT, R-LSTM, DeepFE, R-FC, PIPR, and Topsy-Turvy. The results are given in Table 3.

**Table 3.**
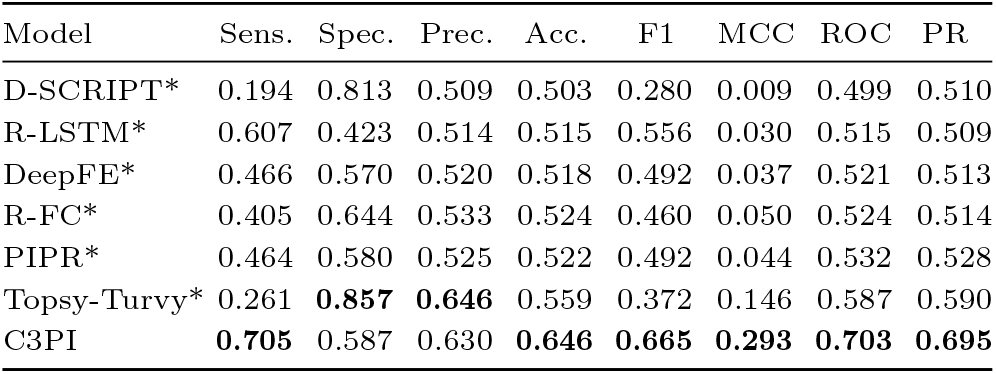
Comparison on the gold standard dataset. The results for methods marked by * are from Bernett *et al*. [5]. The best results are shown in boldface.

| Model | Sens. | Spec. | Prec. | Acc. | F1 | MCC | ROC | PR |
| --- | --- | --- | --- | --- | --- | --- | --- | --- |
| D-SCRIPT* | 0.194 | 0.813 | 0.509 | 0.503 | 0.280 | 0.009 | 0.499 | 0.510 |
| R-LSTM* | 0.607 | 0.423 | 0.514 | 0.515 | 0.556 | 0.030 | 0.515 | 0.509 |
| DeepFE* | 0.466 | 0.570 | 0.520 | 0.518 | 0.492 | 0.037 | 0.521 | 0.513 |
| R-FC* | 0.405 | 0.644 | 0.533 | 0.524 | 0.460 | 0.050 | 0.524 | 0.514 |
| PIPR* | 0.464 | 0.580 | 0.525 | 0.522 | 0.492 | 0.044 | 0.532 | 0.528 |
| Topsy-Turvy* | 0.261 | **0.857** | **0.646** | 0.559 | 0.372 | 0.146 | 0.587 | 0.590 |
| C3PI | **0.705** | 0.587 | 0.630 | **0.646** | **0.665** | **0.293** | **0.703** | **0.695** |

C3PI is by far the winner, with Topsy-Turvy clearly the main competitor. The main conclusion of the study of Bernett *et al*. was that the previous methods are essentially random, with the slight exception of Topsy-Turvy. The composite measures shown visually in Figure 7 indicate this very clearly: accuracy, AUROC, and AUPRC for all methods stay at 50%, then increase slightly for Topsy-Turvy, then increase again, significantly, for C3PI. The trend is most clearly visible for MCC. C3PI’s improvement over Topsy Turvy is 15.54% for accuracy, 79.06% for F1-score, 100.15% for MCC, 19.66% for AUROC and 17.86% for AUPRC.

**Fig. 7.**
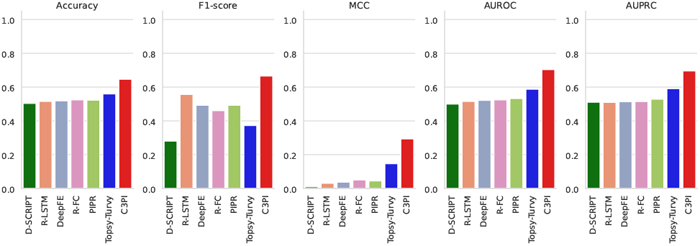
Bar chart visualization of composite parameters – accuracy, F1-score, MCC, AUROC, and AUPRC – from the gold standard dataset comparison (Table 3).

### Ablation study

In order to evaluate to contribution of our puzzler and entangler architecture components, we performed an ablation study where we trained and evaluated the model while removing one or both components on the gold standard dataset. While removing the puzzler component, each protein sequence is padded or truncated to a fixed length of 800 amino acids and passed directly to the model. No permuted sequences are generated. The rest of the architecture remains unchanged, including the dual-branch convolutional encoder, scale-wise entangler, and final classification MLP. For the removal of the entangler component, the layers that fuse protein features at each scale are removed and the six scale-specific embeddings from each protein are concatenated to form a 2016-dimensional feature vector. The classifier is modified accordingly to include the following components:

- Linear layer (2016 → 64) + ReLU + Dropout
- Linear layer (64 → 8) + ReLU + Dropout
- Linear layer (8 → 1) + Sigmoid

Removing both the puzzler and entangler involves performing all the above modifications.

The results of the tests on the gold standard dataset are shown in Table 4. The removal of puzzler or entangler brings a significant drop in performance, more pronounce for the puzzler than for the entangler. Without either one of them, the performance is still above that of Topsy-Turvy. The removal of both components brings the model to essentially random performance, comparable with the other models in Table 3.

**Table 4.** Ablation study. The full model is compared on the gold standard dataset with the model without the entangler component, puzzler component, and both.

| Model | Sens. | Spec. | Prec. | Acc. | F1 | MCC | ROC | PR |
| --- | --- | --- | --- | --- | --- | --- | --- | --- |
| C3PI | 0.705 | 0.587 | 0.630 | 0.646 | 0.665 | 0.293 | 0.703 | 0.695 |
| w/o ent. | 0.648 | 0.561 | 0.596 | 0.604 | 0.621 | 0.209 | 0.646 | 0.634 |
| w/o puz. | 0.569 | 0.618 | 0.598 | 0.593 | 0.583 | 0.187 | 0.569 | 0.634 |
| w/o both | 0.570 | 0.505 | 0.535 | 0.537 | 0.552 | 0.075 | 0.549 | 0.529 |

## Application: NOTCH signalling pathway

The NOTCH signalling pathway is one of the best studied protein interaction pathways [3, 19] and continues to attract interest [6]. The NOTCH pathway is critical in many biomolecular processes, and defects in the pathway are known to be involved in several types of cancers and in several neurological diseases. Compared to other signally pathways, the Notch pathway is comparatively unusual in that it involves a one-on-one interaction between the notch receptor in the plasma membrane and a single ligand.

The interaction data used here has been taken from [11] who used the NOTCH network as a canonical example of networks in biology. This study shows the network, as derived from studies in humans, and includes a total of 50 binary interactions including 33 that involve one of the four NOTCH proteins in humans.

The predictions by D-SCRIPT, Topsy-Turvy and C3PI are used to plot the networks in Figure 8. A summary is given in Table 5. Complete predictions are given in the Supplementary Table 1. D-SCRIPT performs very poorly on this example. C3PI performs clearly better than Topsy-Turvy, as seen from the summary in Table 5 and from Figure 8. The average score for C3PI predictions is 0.644, whereas Topsy-Turvy has 0.490. In spite of significant improvement, plenty of room for improvement remains.

**Table 5.** NOTCH network prediction summary.

| Prediction threshold | D-SCRIPT | Topsy-Turvy | C3PI |
| --- | --- | --- | --- |
| Above 90% | 0 | 0 | 14 |
| Above 75% | 1 | 5 | 21 |
| Above 50% | 1 | 30 | 37 |
| Below 5% | 47 | 4 | 3 |

**Fig. 8.**
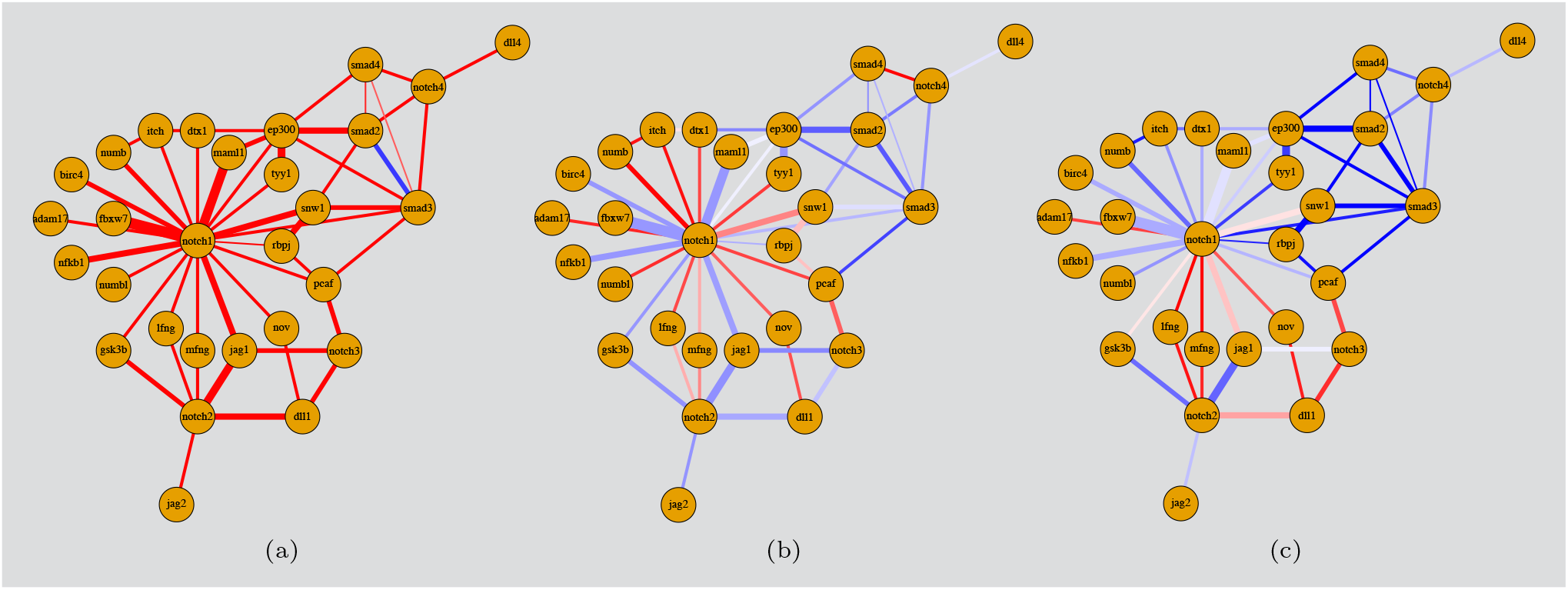
The NOTCH network as predicted by (a) D-SCRIPT, (b) Topsy-Turvy, (c) C3PI. A three-colour scheme is used for the edges: white if the prediction is close to 50%, shades of blue if above 50%, and shades of red if below 50%. (The width of each edge indicates the number of experiments supporting that interaction.)

## Conclusion

We have introduced C3PI, a novel protein-protein interaction prediction model. C3PI uses as input ProtT5 protein embeddings into a complex architecture that includes two novel components, a puzzler and an entangler, which significantly enhance the model’s performance. C3PI clearly outperforms the state-of-the-art methods, being the first PPI prediction method to achieve a significant improvement over random on the leakage-free gold standard dataset of Bernett *et al*. [5]

While the improvements over the state-of-the-art programs are considerable, the tests presented on the golden standard dataset and the NOTCH network indicate that there is still need for improvement in this area.

## Acknowledgements

All our computations were performed on Digital Research Alliance of Canada servers.

## Availability

C3PI is freely available as a web server at c3pi.csd.uwo.ca and as source code from github.com/lucian-ilie/C3PI.

## Author contributions

S.H. designed the architectures, computed the datasets, wrote the software, performed all tests, including installing and running competing methods, and wrote an initial draft of the manuscript. G.B.G. designed and analyzed the NOTCH application and wrote that section. L.I. proposed the problem and designed the project, suggested the use of embeddings, analyzed the results, supervised the work, and wrote the final version of the manuscript.

