## Supplemental file for "C3PI: Component Puzzle Protein-Protein Interaction Prediction"

– Supplementary material –

SeyedMohsen Hosseini, G. Brian Golding, Lucian Ilie*

### 1 Application: NOTCH signalling pathway

Supplementary Table 1 contains the complete predictions by D-SCRIPT, Topsy-Turvy and C3PI for the NOTCH network, which were used to plot the graphs in Figure 8 in the main document, as well as compute the statistics in Table 5.

Supplementary Table 1: D-SCRIPT, Topsy-Turvy and C3PI complete predictions for the NOTCH network.

| Protein1 | Protein2 | Links | D-Script | Topsy-Turvy | C3PI | Protein1 | Protein2 | Links | D-Script | Topsy-Turvy | C3PI |
|---|---|---|---|---|---|---|---|---|---|---|---|
| adam17 | notch1 | 2 | 0.004 | 0.074 | 0.128 | notch1 | nov | 2 | 0.004 | 0.182 | 0.175 |
| dll1 | notch2 | 4 | 0.004 | 0.671 | 0.321 | notch1 | numb | 3 | 0.004 | 0.007 | 0.796 |
| dll1 | notch3 | 3 | 0.004 | 0.619 | 0.091 | notch1 | numbl | 2 | 0.004 | 0.088 | 0.708 |
| dll1 | nov | 2 | 0.004 | 0.158 | 0.057 | notch1 | pcaf | 2 | 0.004 | 0.132 | 0.644 |
| dll4 | notch4 | 2 | 0.004 | 0.551 | 0.640 | notch1 | rbpj | 1 | 0.004 | 0.654 | 0.929 |
| ep300 | dtx1 | 2 | 0.004 | 0.735 | 0.665 | notch1 | smad3 | 2 | 0.004 | 0.640 | 0.940 |
| ep300 | maml1 | 3 | 0.004 | 0.527 | 0.557 | notch1 | snw1 | 4 | 0.004 | 0.260 | 0.445 |
| ep300 | tyy1 | 5 | 0.006 | 0.707 | 0.880 | notch1 | tyy1 | 2 | 0.004 | 0.115 | 0.880 |
| itch | dtx1 | 2 | 0.004 | 0.449 | 0.776 | notch2 | gsk3b | 3 | 0.004 | 0.713 | 0.791 |
| jag1 | notch1 | 4 | 0.004 | 0.690 | 0.383 | notch3 | pcaf | 3 | 0.004 | 0.187 | 0.142 |
| jag1 | notch2 | 5 | 0.004 | 0.718 | 0.802 | notch4 | smad2 | 2 | 0.004 | 0.758 | 0.747 |
| jag1 | notch3 | 3 | 0.004 | 0.730 | 0.530 | notch4 | smad3 | 2 | 0.004 | 0.705 | 0.725 |
| jag2 | notch2 | 2 | 0.004 | 0.709 | 0.623 | notch4 | smad4 | 2 | 0.004 | 0.008 | 0.783 |
| lfng | notch1 | 2 | 0.004 | 0.144 | 0.004 | numb | itch | 2 | 0.004 | 0.037 | 0.944 |
| lfng | notch2 | 2 | 0.004 | 0.344 | 0.039 | pcaf | rbpj | 2 | 0.004 | 0.378 | 0.971 |
| mfng | notch1 | 2 | 0.004 | 0.339 | 0.007 | rbpj | snw1 | 4 | 0.004 | 0.373 | 0.996 |
| mfng | notch2 | 2 | 0.004 | 0.270 | 0.078 | smad2 | ep300 | 4 | 0.004 | 0.818 | 0.999 |
| notch1 | birc4 | 3 | 0.004 | 0.701 | 0.665 | smad2 | smad3 | 3 | 0.889 | 0.826 | 1.000 |
| notch1 | dtx1 | 2 | 0.004 | 0.137 | 0.665 | smad2 | smad4 | 1 | 0.077 | 0.717 | 1.000 |
| notch1 | ep300 | 2 | 0.004 | 0.527 | 0.600 | smad2 | snw1 | 2 | 0.004 | 0.678 | 1.000 |
| notch1 | fbxw7 | 7 | 0.004 | 0.701 | 0.665 | smad3 | ep300 | 2 | 0.004 | 0.779 | 0.998 |
| notch1 | gsk3b | 2 | 0.004 | 0.701 | 0.450 | smad3 | pcaf | 2 | 0.004 | 0.870 | 1.000 |
| notch1 | itch | 2 | 0.004 | 0.030 | 0.716 | smad3 | smad4 | 1 | 0.182 | 0.651 | 1.000 |
| notch1 | maml1 | 6 | 0.004 | 0.701 | 0.557 | smad3 | snw1 | 3 | 0.004 | 0.560 | 1.000 |
| notch1 | nfkb1 | 4 | 0.004 | 0.701 | 0.665 | smad4 | ep300 | 2 | 0.004 | 0.708 | 0.995 |
